# 6-Phosphogluconolactonase is critical for the efficient functioning of the pentose phosphate pathway

**DOI:** 10.1101/2023.11.29.569206

**Authors:** Léa Phégnon, Julien Pérochon, Sandrine Uttenweiler-Joseph, Edern Cahoreau, Pierre Millard, Fabien Létisse

**Affiliations:** Toulouse Biotechnology Institute, Université de Toulouse, INSA, UPS, Toulouse, France; MetaToul-MetaboHUB, National Infrastructure of Metabolomics and Fluxomics, Toulouse, France; Institut de Pharmacologie et de Biologie Structurale (IPBS), Université de Toulouse, CNRS, Université Toulouse III - Paul Sabatier (UT3), Toulouse, France

**Author notes:** **Corresponding author** Fabien Létisse, IPBS-Toulouse, BP64182, 205 route de Narbonne, 31077 Toulouse Cedex04, https://www.ipbs.fr. Co first authors (Léa Phégnon, Julien Pérochon).

**Keywords:** Metabolic network, Metabolic fluxes, Metabolic bypass, phosphogluconolactone, gluconolactone, ^13^C NMR, ^13^P NMR, microorganisms

## Abstract

The metabolic networks of microorganisms are remarkably robust to genetic and environmental perturbations. This robustness stems from redundancies such as gene duplications, isoenzymes, alternative metabolic pathways, and also from non-enzymatic reactions. In the oxidative branch of the pentose-phosphate pathway (oxPPP), 6-phosphogluconolactone hydrolysis into 6-phosphogluconate is catalysed by 6-phosphogluconolactonase (Pgl) but in the absence of the latter, the oxPPP flux is thought to be maintained by spontaneous hydrolysis. However, in Δ*pgl Escherichia coli*, an extracellular pathway can also contribute to pentose-phosphate synthesis. This raises question as to whether the intracellular non-enzymatic reaction can compensate for the absence of 6-phosphogluconolactonase and, ultimately, on the role of 6-phosphogluconolactonase in central metabolism. Our results validate that the bypass pathway is active in the absence of Pgl, specifically involving the extracellular spontaneous hydrolysis of gluconolactones to gluconate. Under these conditions, metabolic flux analysis reveals that this bypass pathway accounts for the entire flux into the oxPPP. This alternative metabolic route - partially extracellular - sustains the flux through the oxPPP necessary for cell growth, albeit at a reduced rate in the absence of Pgl. Importantly, these findings imply that intracellular non-enzymatic hydrolysis of 6-phosphogluconolactone does not compensate for the absence of Pgl. This underscores the crucial role of Pgl in ensuring the efficient functioning of the oxPPP.

## INTRODUCTION

Metabolic networks are a set of interconnected chemical reactions, almost all of which are catalysed by enzymes. The interplay between chemical reactions provides alternative routes for adaptation to genetic or environmental perturbations. The network organisation of metabolic systems thus underpins the metabolic robustness and adaptability of cells. The central metabolism of *Escherichia coli* is a model of metabolic robustness since very few of the associated genes are indispensable for growth on glucose minimal medium [1,2]. Furthermore, *E. coli* knock-out mutants lacking key central metabolic enzymes have similar growth phenotypes [3–7]. This robustness is in part the result of metabolic flux rerouting [6,7] but stems also from local compensatory mechanisms based on redundancy, such as the presence of isozymes and alternative pathways [2–5,7].

Compensatory mechanisms involving non-enzymatic reactions are harder to identify because they cannot be studied by metabolic reconstruction using comparative genomics [8]. These mechanisms are based on specific or non-specific chemical reactions that occur either exclusively non-enzymatically within the metabolic network, or in parallel to existing enzyme functions [9]. The second step in the oxidative branch of the pentose-phosphate pathway (oxPPP), the spontaneous hydrolysis of δ-6-phosphogluconolactone (δ-6PGL) into 6-phosphogluconate (6PGNT) is an archetypal example of a reaction that can occur enzymatically, catalysed by 6-phosphogluconolactonase (Pgl, EC 3.1.1.31), and non-enzymatically, without Pgl activity. However, Pgl’s main role, rather than maintaining the flux through the oxPPP, is thought to be preventing the formation of unwanted side-products by covalent modification with highly reactive 6PGLs (δ (1-5) and γ (1-4)) [10–12]. In pioneering studies, Kupor and Fraenkel [13,14] found that absence of the *pgl* gene indeed slowed the growth of *E. coli* on glucose, but more importantly perhaps, they also discovered an alternative pathway bypassing Pgl, involving dephosphorylation and secretion of gluconolactone, spontaneous abiotic hydrolysis in the medium, and re-import of gluconate and phosphorylation [14]. This “Pgl bypass” can therefore provide the same anabolic precursors (NADPH and pentose-phosphates) as the canonical oxPPP.

In *E. coli*, one fifth of glucose uptake is directed towards the oxPPP to provide the required anabolic precursors [6], highlighting the importance of this pathway. In the absence of Pgl, it is thought that this flux is maintained by rapid non-enzymatic hydrolysis of δ-6PGL rather than by the Pgl bypass [7,11,14,15]. However, 6PGLs have a non-negligible lifetime [11], raising the questions whether spontaneous δ-6PGL hydrolysis is fast enough to maintain a high flux through the oxPPP in the absence of Pgl and how the bypass pathway might contribute.

In this study, we used a metabolic flux approach to investigate *E. coli* oxPPP function in the absence of Pgl to (i) reinvestigate the extracellular bypass pathway identified by Kupor and Fraenkel using state-of-the-art methods, (ii) quantify its contribution to oxPPP flux, and (iii) elucidate the metabolic function of Pgl. In the absence of Pgl, we found that virtually all carbon flux through the oxPPP was channelled through the Pgl bypass, and that the contribution of non-enzymatic δ-6PGL hydrolysis was negligible. This in turn suggests that the catalytic function of Pgl is required to maintain high flux through the canonical oxPPP, and that the metabolic role of this enzyme is more substantial than simple house-cleaning duties.

## RESULTS

### Slow growth in *E. coli* Δ*pgl* is accompanied by gluconate accumulation in the culture medium

We first sought to determine the growth phenotype of an *E. coli* K-12 strain lacking *pgl*, grown on minimal medium with glucose (15 mM) as sole carbon source (Table 1 and Figure 1). In agreement with Kupor and Fraenkel’s [14] findings, the Δ*pgl* strain grew much slower than the WT. The specific rate of acetate production was markedly increased in the Δ*pgl* strain, resulting in a higher acetate yield (0.54 ± 0.03 mol/mol) than in the WT strain (0.31 ± 0.02 mol/mol), as observed previously [7]. In addition to acetate and other metabolites previously detected by ^1^H NMR [16], significant amounts of gluconate were also detected, but only in the culture medium of the Δ*pgl* strain (Figure 2A-C). Gluconate accumulation represented approximately 2 % of the total carbon flux entering the central metabolic pathways of the *Δpgl* strain (Table 1). Note that no 6PGNT was detected in the culture supernatant of the *Δpgl* strain (Figure 2B, D). Complementation of the Δ*pgl* strain with the WT gene restored the growth rate determined for the WT *E. coli*, without any gluconate accumulation (Table 1), demonstrating that reduced growth and gluconate excretion are related to the absence of Pgl.

**Table 1:** Growth parameters of the different *E. coli* BW25113 strains grown in minimal media with 15 mM glucose. The growth of the *E. coli* BW25113 *Δpgl* strain was compared to the WT strain’s and that of the complemented strain, Δ*pg*l::pZA23*pgl*, obtained by transformation of the *Δpg*l strain with pZA23::*pgl* plasmid and of the Δ*pgl*::pZA23 strain with an empty plasmid. *µ*_max_, growth rate; q_Glc_, specific glucose uptake rate; q_Ac_, specific net acetate production rate; q_Gnt_, specific gluconate accumulation rate. Results are the mean ± one SD of three biologically independent samples.

| Parameters | WT | $\Delta pgl$ | $\Delta pgl::pZA23pgl$ | $\Delta pgl::pZA23$ |
| --- | --- | --- | --- | --- |
| $\mu_{max} (h^{-1})$ | $0.61 \pm 0.01$ | $0.43 \pm 0.01$ | $0.57 \pm 0.01$ | $0.41 \pm 0.01$ |
| $q_{Glc} (mmol \cdot [g_{CDW} \cdot h]^{-1})$ | $7.77 \pm 0.17$ | $8.52 \pm 0.39$ | $10.11 \pm 0.83$ | $7.60 \pm 0.08$ |
| $q_{Ac} (mmol \cdot [g_{CDW} \cdot h]^{-1})$ | $2.37 \pm 0.15$ | $4.56 \pm 0.16$ | $7.60 \pm 0.31$ | $3.95 \pm 0.16$ |
| $q_{Gnt} (mmol \cdot [g_{CDW} \cdot h]^{-1})$ | 0 | $0.17 \pm 0.01$ | 0 | $0.14 \pm 0.02$ |

**Figure 1.**
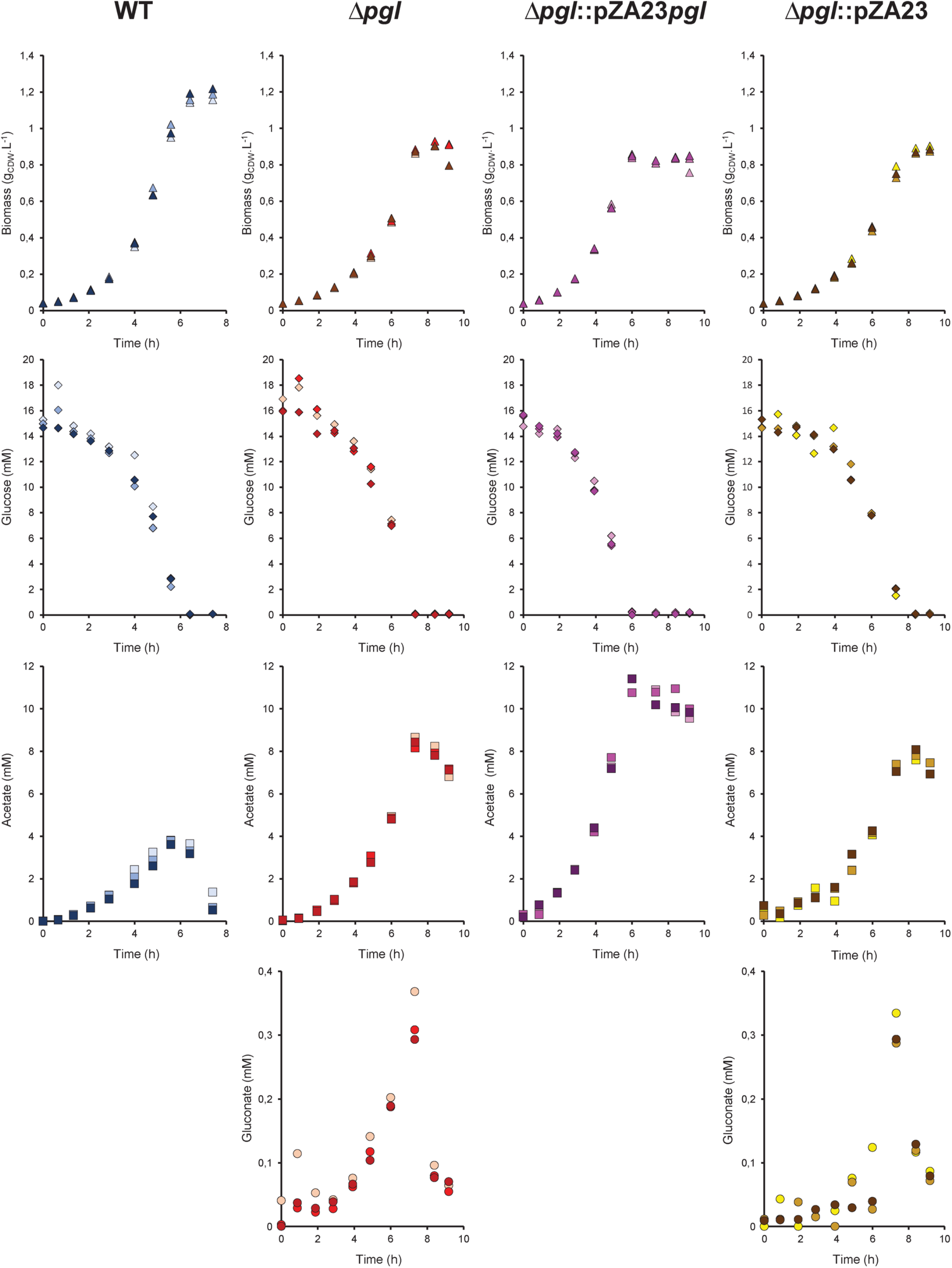
Growth phenotype of *E. coli* BW25113 WT (blue), Δ*pgl* (red), Δ*pgl*::pZA23*pgl* complement (purple) and Δ*pgl*::pZA23 control (yellow) strains cultivated in M9-based medium supplemented with glucose (pH 7). Extracellular metabolite concentrations were quantified by ^1^H-NMR in culture supernatant. Gluconate was only detected in the culture medium of the Δ*pgl* and Δ*pgl*::pZA23 strains. n=3 biological replicates, colour gradient.

**Figure 2.**
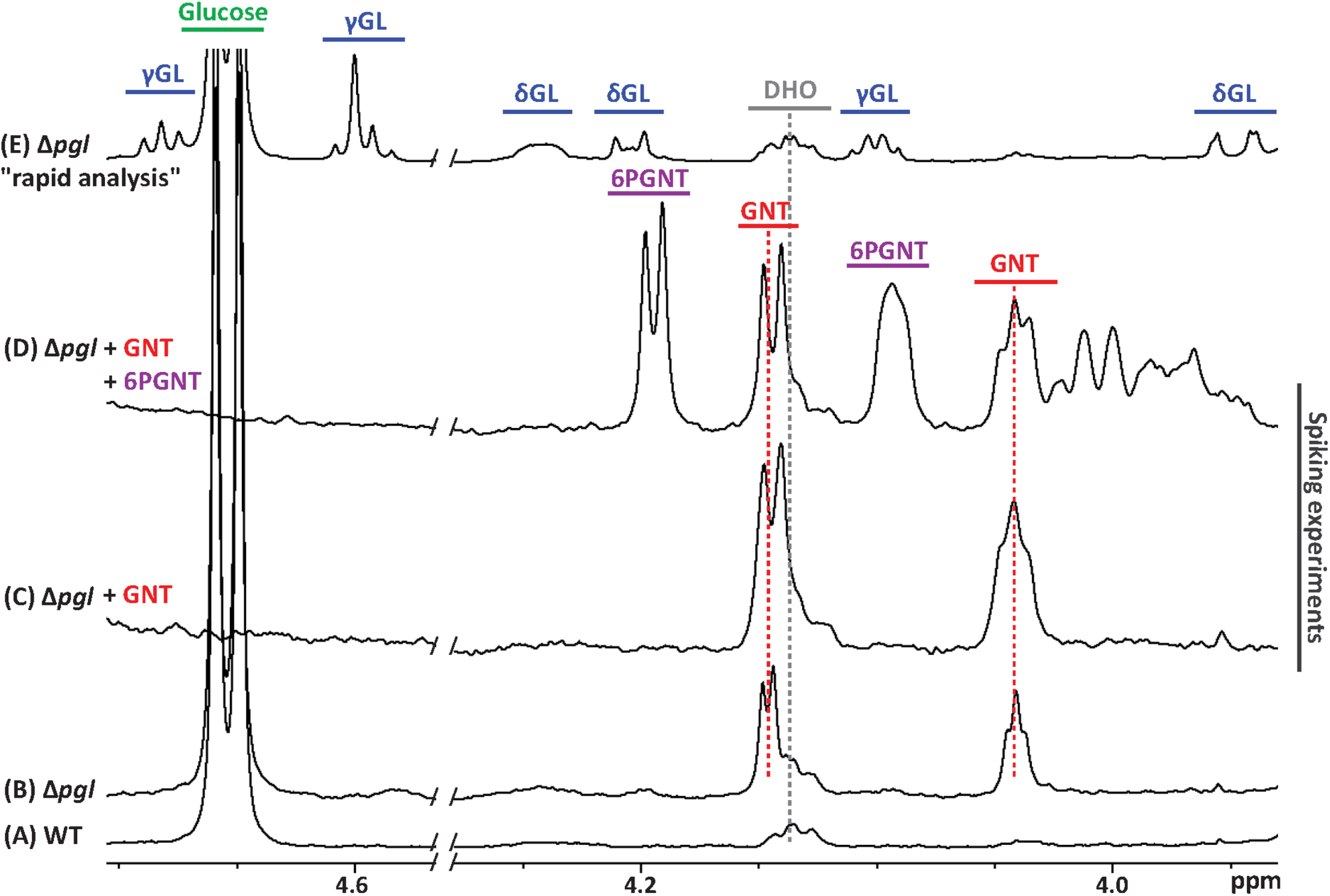
Gluconate, δ- and γ-gluconolactone identification by ^1^H NMR analysis of *E. coli* BW25113 Δ*pgl* culture supernatants. Selected region of the 1D ^1^H NMR spectrum of the culture supernatant of WT *E. coli* BW25113 **(A)** (no detectable gluconate (GNT)) and *E. coli* BW25113 Δ*pgl* **(B)** (estimated GNT concentration: 0.35 mM). Culture broth was centrifugated at 12,000 g for 3 min and supernatant was stored at −20 °C prior to NMR analysis. The so-obtained culture supernatant of the Δ*pgl* strain was then spiked with 0.2mM GNT **(C)**, and with 0.2 mM GNT and 0.7 mM of 6-phosphogluconate (6PGNT) **(D)**. Selected region of the 1D ^1^H NMR spectrum of the culture supernatant of Δ*pgl* strain using a “rapid analysis” procedure **(E)**. Culture broth was filtered on a polyethersulfone membrane with a 0.2 µm pore size and placed immediately on ice before NMR analysis. NMR spectra were recorded approximately 5 min after sample withdrawal. Both isomeric forms of GL (δ and γ) were detected but GNT was beneath the detection limit. DHO: Dihydroorotate.

### Gluconate is formed by abiotic degradation of gluconolactone

In the extracellular Pgl bypass proposed by Kupor and Fraenkel [14], gluconolactones are hydrolysed into gluconate abiotically in the culture medium. The obvious explanation for the presence of gluconate in the extracellular medium is therefore that gluconolactones are fully converted into gluconate within the time taken to sample the culture medium and store the samples. To prevent abiotic hydrolysis, the culture medium was sampled by filtration, placed immediately on ice and then analysed by NMR, with the first spectrum recorded approximately 5 min after sample withdrawal. Both isomeric forms of gluconolactone (δ and γ) were detected in equal amounts whereas the gluconate concentration was below the detection limit (Figure 2E). 6PGL was not detected either. Importantly, the total amount of gluconolactones was similar to the amount of gluconate detected in the corresponding sample obtained with the initial procedure, indicating that rapidly reducing the sample temperature prevents the spontaneous degradation of gluconolactones. To confirm this effect, we measured the rate constants of gluconolactone spontaneous non-enzymatic hydrolysis in fresh minimal medium containing commercial gluconolactone using real-time ^1^H-NMR (Figure 3A). The degradation constants obtained by fitting the gluconolactone concentration–time curves (Figure 3C, D) assuming a first-order process [17] were 0.36 ± 0.01 h^−1^ at 7 °C and 2.58 ± 0.02 h^−1^ at 37 °C (Figure 3F). At 7 °C therefore, the amount of gluconolactone hydrolysed during 5 min of sample preparation is negligible. Using the same approach, the hydrolysis rate of δ-6PGL was found to be 3.97 ± 0.04 h^−1^ at 37 °C, about 6-7 times higher than at 5 °C [11] (Figure 3B, E, G) and the same order of magnitude as measured for gluconolactone.

**Figure 3.**
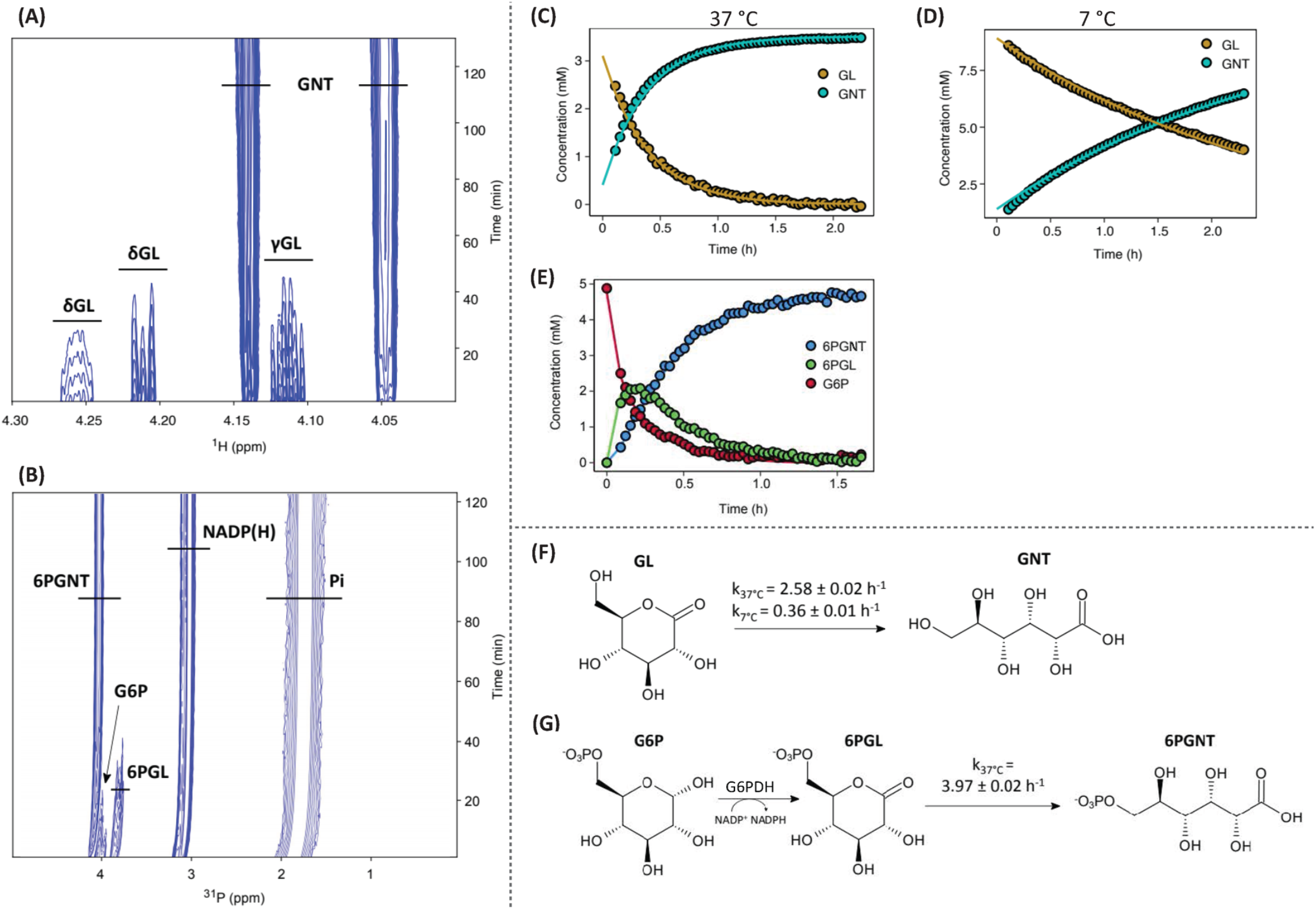
Determination of spontaneous hydrolysis rate constants of gluconolactone (GL) and 6-phosphogluconolactone (6PGL). Pseudo-2D ^1^H spectra at 37 °C, pH 7.2 in M9 synthetic minimal media of spontaneous gluconolactone **(A)** and 6-phosphogluconolactone **(B)** hydrolysis; T0 corresponds to the beginning of NMR acquisition (6 min 35 s and 5 min 25 s after gluconolactone and 6-phosphogluconolactone addition respectively). Concentrations extracted from pseudo-2D spectra (dots) and fitted by COPASI model (lines) for GL and GNT at 37 °C **(C)** and 7 °C **(D)**, and for 6PGNT, G6P and 6PGL **(E)**. Reaction schemes for the spontaneous hydrolysis of GL into GNT **(F)** and for enzymatic production of 6PGL from G6P *via* G6PDH followed by spontaneous 6PGL hydrolysis into 6PGNT **(G)** with the values of the hydrolysis rate constant (k) so-determined.

The detection of gluconate in the culture medium in the above experiments is therefore an experimental artefact arising from the hydrolysis of gluconolactones during the sampling process. The Δ*pgl* strain produces gluconolactone (not gluconate) during growth on glucose.

### Glucose-derived gluconate is metabolized by canonical gluconate metabolism

Under growth conditions at 37 °C, gluconolactones in the culture medium spontaneously hydrolyse to gluconate. However, gluconate remained below the limit of detection when the culture medium was sampled by filtration and rapidly cooled. Therefore, we hypothesized that the cells’ flux capacity to consume gluconate is high enough to metabolize all the gluconate produced by spontaneous hydrolysis. To confirm this, we blocked gluconate utilization by knocking out the two gluconokinases, GntK and IdnK, known to activate gluconate after uptake [18,19]. As reported previously [20], the Δ*gntK* Δ*idnK* double mutant did not grow on D-gluconate as sole carbon source. As expected therefore, gluconate accumulation was observed when the Δ*pgl*Δ*gntK*Δ*idnK* triple mutant was grown on glucose, with a specific extracellular accumulation rate of 1.13 ± 0.04 mmol·(g_CDW_·h)^−1^ (Figure 4). These results are consistent with the Pgl bypass topology described by Kupor and Fraenkel [14] (Figure 5).

**Figure 4.**
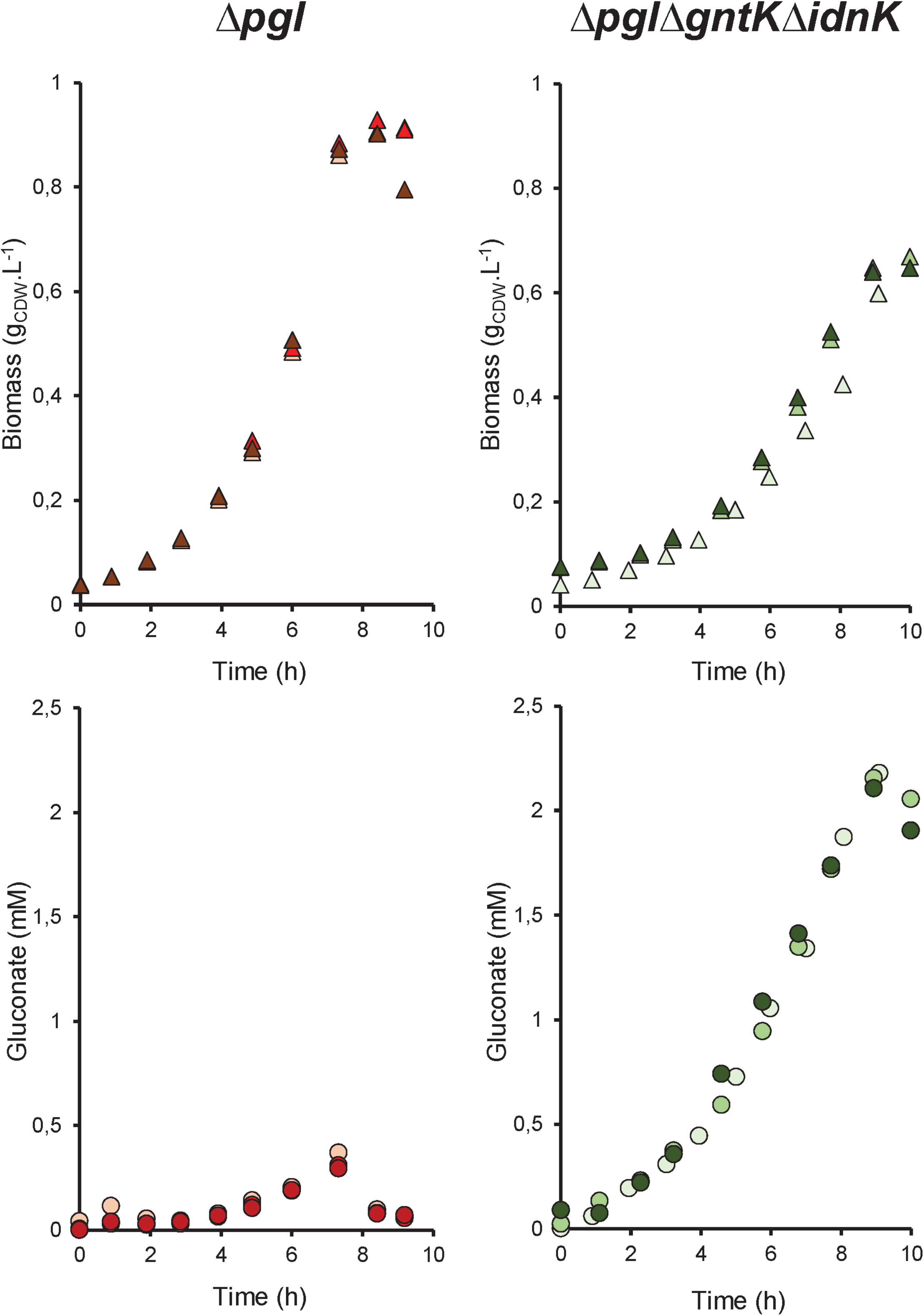
Gluconate accumulation in the culture medium of *E. coli* BW25113 Δ*pgl* (red) and *E. coli* BW25113 Δ*pgl*Δ*gntK*Δ*idnK* (green) strains cultivated in M9-based medium supplemented with glucose (pH 7). Extracellular gluconate concentration was quantified by ^1^H-NMR in culture supernatant. n=3 biological replicates, colour gradient.

**Figure 5.**
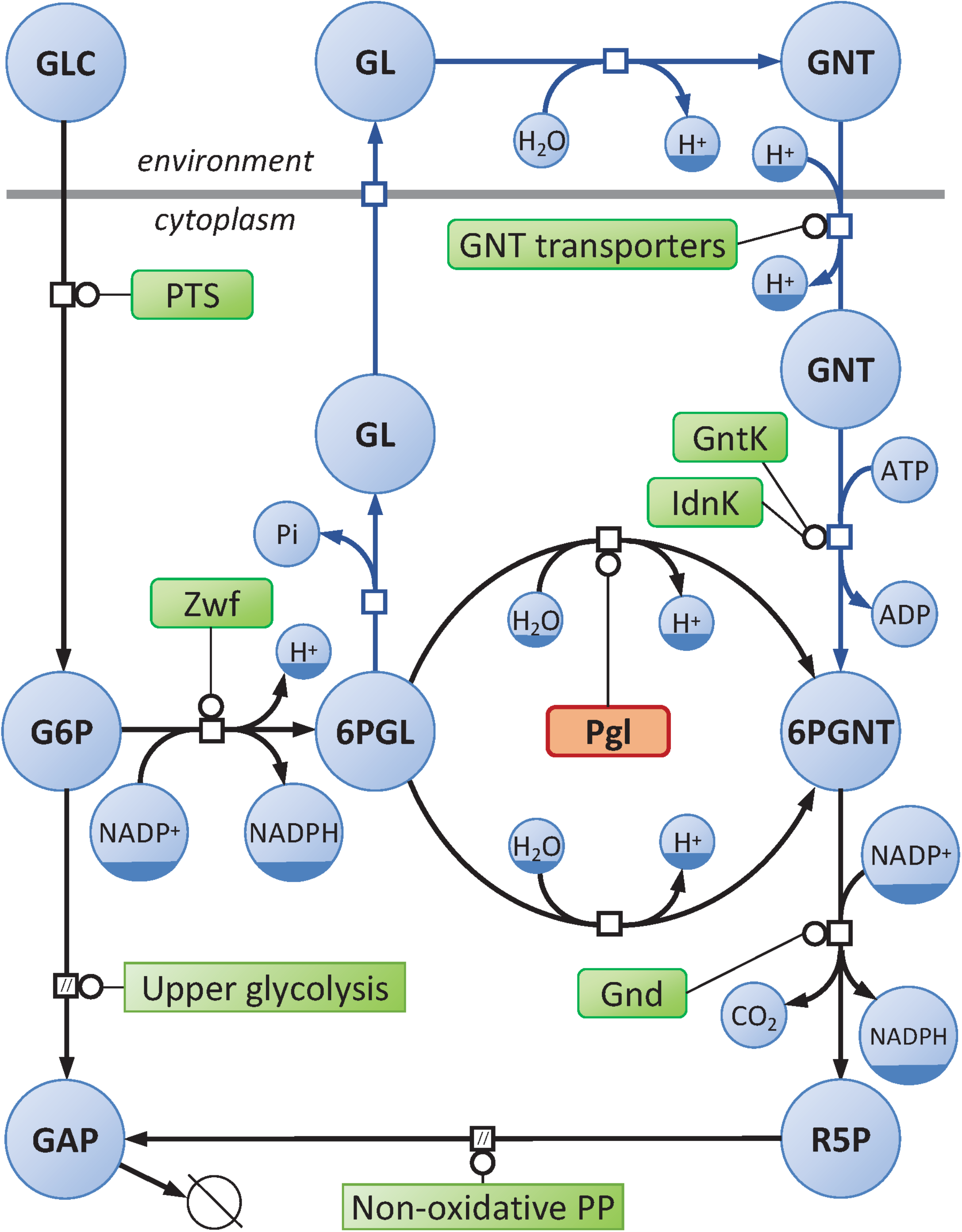
Schematic representation of the Pgl bypass and of the oxidative branch of the pentose-phosphate (PP) pathway in *E. coli*, in Systems Biology Graphical Notation format [46] (https://sbgn.github.io). Circles represent metabolites and rounded rectangles represent enzymes. GLC, glucose; G6P, glucose-6-phosphate; GAP, glyceraldehyde-3-phosphate; 6PGL, 6-phosphogluconolactone; GL, gluconolactone; GNT, gluconate; 6PGNT, 6-phosphogluconate; R5P, ribulose-5-phosphate; PTS, phosphoenolpyruvate:glucose phosphotransferase system; Zwf, glucose-6-phosphate dehydrogenase; GntK, gluconate kinase; IdnK, thermosensitive gluconate kinase; Gnd, 6-phosphogluconate dehydrogenase.

An unforeseen consequence of gluconokinase deletion was that the Δ*pgl*Δ*gntK*Δ*idnK* triple mutant had a much slower growth rate than the Δ*pgl* strain (0.29 ± 0.01 h^-1^ *vs*. 0.43 ± 0.01 h^-1^). This highlights the importance of metabolic activity in the bypass pathway to sustain growth. To investigate this further, we quantified the metabolic flux through the extracellular bypass using ^13^C metabolic flux analysis. Since the bypass pathway and the non-enzymatic hydrolysis of δ-6PGL operate in parallel without carbon scrambling, their relative contributions to the oxPPP flux cannot be determined using stationary ^13^C labelled experiments alone. We therefore investigated the contributions of both routes in the Δ*pgl* strain using stationary and non-stationary metabolic flux analyses.

### Deletion of *pgl* barely modifies the contribution of the oxidative branch of the PPP to glucose metabolism

We first quantified the distribution of glucose-6-phosphate between the Embden-Meyerhof-Parnas (EMP) pathway and the oxPPP in the Δ*pgl* strain and the WT strain (as control) grown on minimal M9 medium supplemented with [1-^13^C]-glucose as sole carbon source. Metabolic fluxes were inferred from quantitative measurements of ^13^C incorporation into alanine *via* pyruvate (Figure 6). In keeping with previous results [6,21], 17 ± 1 % of glucose was channelled through the oxPPP in the WT strain. In the Δ*pgl* strain, the oxPPP accounted for 15 ± 1 % of glucose uptake, a similar contribution as in the WT strain. This suggests that the absence of Pgl does not lead to a rewiring of central metabolic fluxes, in contrast to what has been observed in the absence of glucose-6-phosphate dehydrogenase, the first enzyme in the oxPPP [6,7]. In the *Δpgl* strain, the metabolic flux through the oxPPP must therefore be maintained by the Pgl bypass and/or by non-enzymatic intracellular hydrolysis of δ-6PGL.

**Figure 6.**
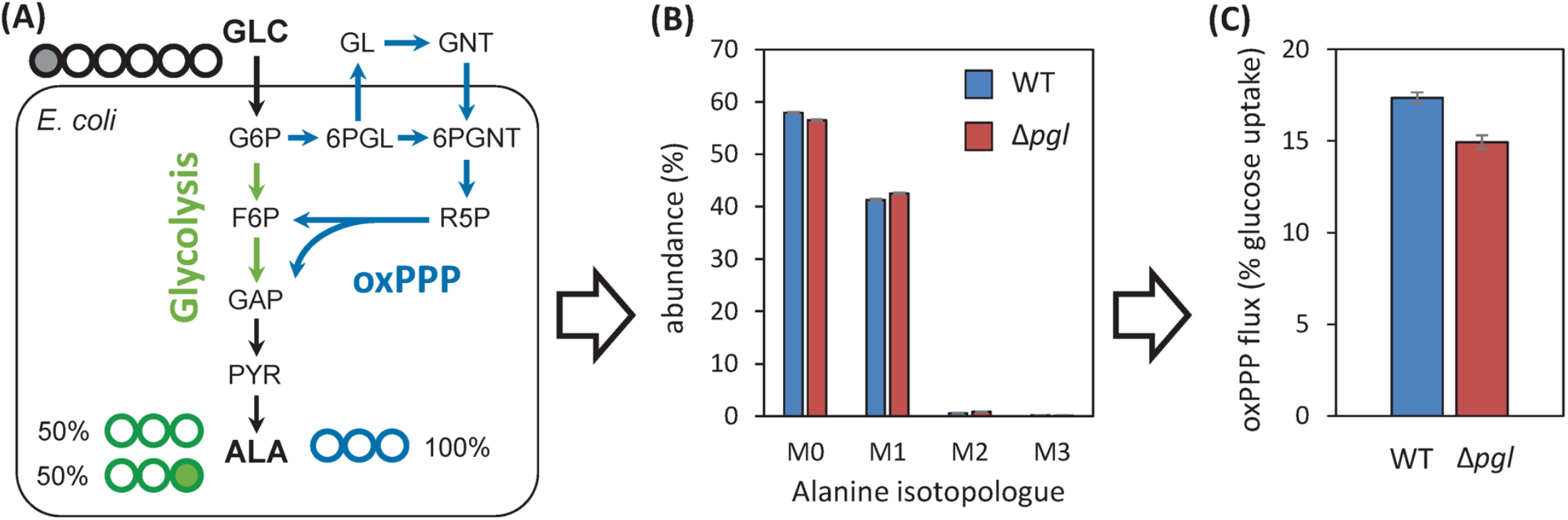
Metabolic fluxes analysis of *E. coli* BW25113 wild-type (WT) and Δ*pgl* strains. **(A)** Cells were grown on 1-^13^C-glucose, which can be metabolized through glycolysis (green pathway) and the pentose phosphate pathway (blue pathway). Filled (empty) circles represent ^13^C (^12^C) atoms. **(B)** Isotopologue distribution of alanine measured by mass spectrometry. The different alanine isotopologues are denoted as M0, M1, M2 and M3, indicating the number of ^13^C atoms present in the molecule. The results are presented as mean ± one SD (below 1 % for all isotopologues) (n=3 biological replicates). **(C)** The partitioning of carbon fluxes between glycolysis and the pentose phosphate pathway was quantified from the data shown in panel B. The bars depict the mean value of the relative contributions of glycolysis and oxPPP, calculated from three independent biological replicates (n=3), with the error bars representing ± one standard deviation from the mean.

### In the absence of Pgl, oxPPP flux passes exclusively though the Pgl bypass

To quantify the contributions of the extracellular bypass and of intracellular non-enzymatic δ-6PGL hydrolysis to oxPPP flux, we performed a non-stationary carbon labelling experiment wherein ^13^C labelled glucose was added at the mid-exponential phase to *E. coli* Δ*pgl* growing on unlabelled glucose (Figure 7A, C). We used [2-^13^C]-glucose because the NMR signals corresponding to [2-^13^C]-glucose and [2-^13^C]-gluconolactones formed from [2-^13^C]-glucose, are well resolved in 1D,^1^H NMR spectra, allowing accurate quantification of the total concentrations of unlabelled and labelled gluconolactone (Figure 7B) over time, sampling the culture medium every 30 min after ^13^C labelled glucose was added. Metabolic fluxes were calculated using a differential equation model of the metabolic network shown in Figure 7D. The flux through the oxPPP and the constant rate of gluconolactone hydrolysis were set to their experimentally determined values (Figure 3). The metabolic fluxes were then obtained by fitting the time-course concentrations of biomass, (unlabelled and labelled) glucose and (total and ^13^C-labelled) gluconolactone. Sensitivity analysis confirmed that the system was identifiable based on the available data, meaning that all fluxes were determined with high precision (relative standard deviation below 5 % in each of the three independent biological replicates).

**Figure 7.**
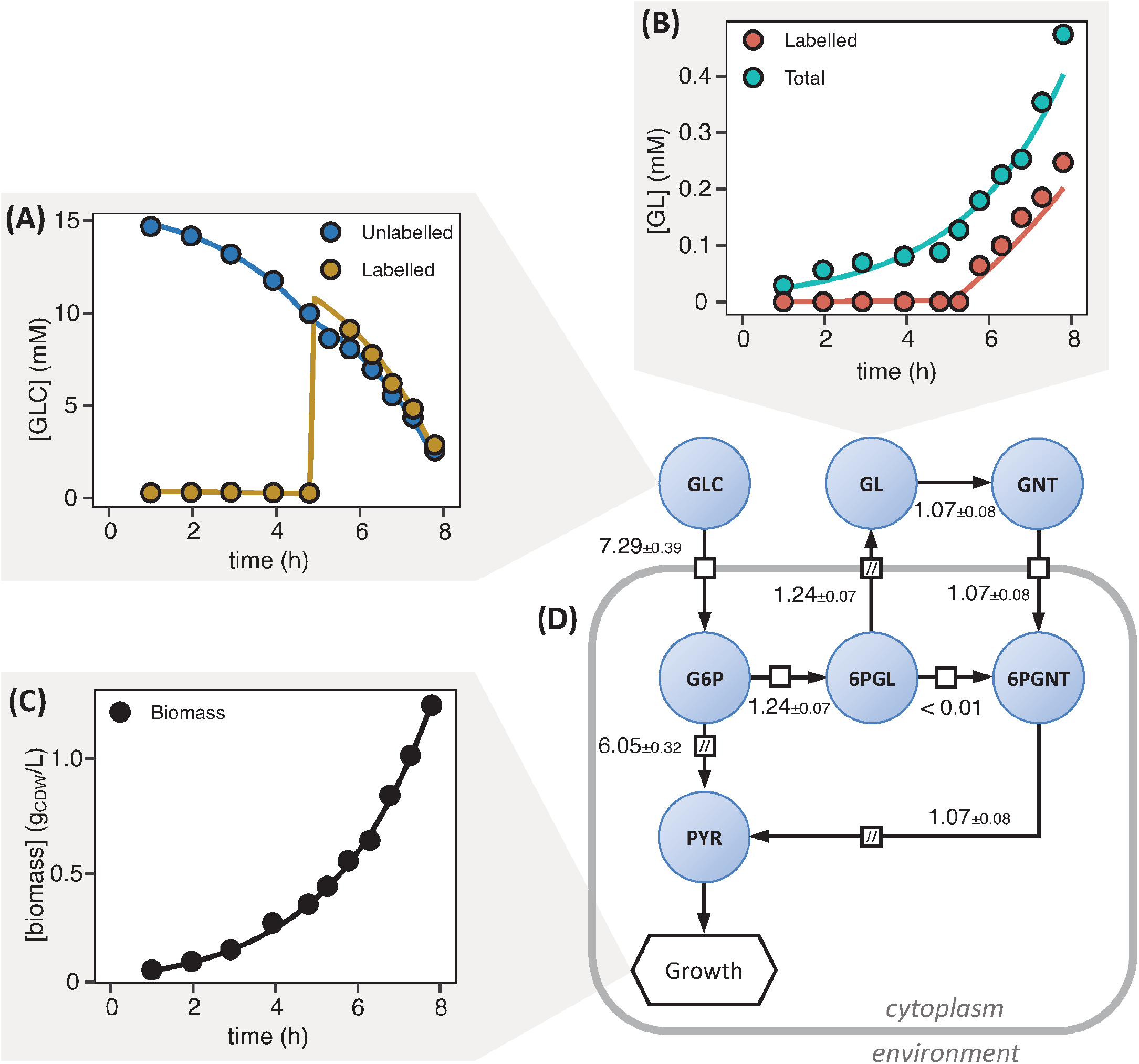
Fluxes in the oxPPP of *E. coli* BW25113 Δ*pgl* quantified by non-stationary ^13^C-metabolic flux analysis. Concentrations of labelled and unlabelled glucose **(A)**. [2-^13^C]-glucose was added at mid-exponential growth (approx. 5 h). Gluconolactone (labelled and total) concentrations **(B)**. Growth of the Δ*pgl* strain as a function of time **(C)**. Metabolic network of the model used to calculate fluxes involved in the Pgl bypass, with absolute flux results **(D)**. Values are the means calculated from three independent biological replicates (n=3) ± one standard deviation.

The estimated glucose uptake rates and the growth rates were consistent with data obtained from unlabelled experiments (Table 1). The intracellular branch accounts for a negligible proportion of oxPPP flux (< 1 ± 1 % of glucose uptake), which is fully channelled through the extracellular bypass (99 ± 1 %). The bypass carries a flux of 1.07 ± 0.08 mmol·(g_CDW_·h)^−1^ which is similar to the flux through the canonical oxPPP in the WT strain and much higher than the gluconolactone accumulation flux in unlabelled experiments (0.17 mmol·(g_CDW_.h)^−1^, Table 1). This flux value is also consistent with the specific gluconate accumulation rate measured in the Δ*pgl*Δ*gntK*Δ*idnK* triple mutant (1.13 ± 0.04 mmol·(g_CDW_·h)^−1^). Overall, these results demonstrate the absence of intracellular flux through the canonical oxPPP under these circumstances, and reveal the key contribution of the extracellular Pgl bypass to R5P biosynthesis in the Δ*pgl* strain. These results also explain the severe growth restriction observed when the Pgl bypass is blocked by the deletion of gluconokinases.

## DISCUSSION

To our knowledge this work is the first time the metabolic activity of the Pgl bypass has been quantified, despite its discovery in *E. coli* more than fifty years ago [14]. Our NMR results confirm the bypass topology proposed by Kupor and Fraenkel, although the 6PGL dephosphorylation and (P-)gluconolactone secretion steps remain to be elucidated. Metabolite phosphatases are generally cytosolic, although some have been shown to be periplasmic, notably those involved in nucleotide salvage [22]. If 6PGL is secreted in its phosphorylated form, it should be found in the culture medium based on the spontaneous hydrolysis rate estimated in this work. Since 6PGL was not detected, this suggests it is dephosphorylated intracellularly and that it is the resulting gluconolactone that is then secreted. Nevertheless, further studies are required to identify the genes involved in dephosphorylation and secretion of gluconolactones and to unequivocally resolve the first two steps of the Pgl bypass.

Because of this bypass, absence of Pgl leads to substantial gluconolactone excretion – about 17 % of the total carbon flux entering the cell. This process, which resembles directed overflow mechanism [23], prevents 6PGL intracellular accumulation, avoiding covalent modifications of proteins [24] and other nucleophiles [10] in the presence of these highly reactive electrophiles. Gluconolactone excretion may therefore be a cleansing mechanism that avoids the formation of toxic metabolites, the function generally ascribed to Pgl [12]. The Pgl bypass may therefore have arisen fortuitously because the unstable gluconolactones excreted into the environment spontaneously hydrolyse into gluconate, one of *E. coli*’s preferred carbon sources [25].

The Pgl bypass produces pentose-phosphates and NAPDH from G6P with the same stoichiometry as the canonical oxPPP. However, this extracellular metabolic route has a significant energetic cost for the cell. First, one extra mole of ATP is required for the phosphorylation of gluconate by gluconokinases (GntK and IdnK). Second, gluconate import is mediated by proton symport systems driven by the proton motive force (with a stoichiometry of one proton per gluconate [26,27]), thereby consuming protons that could otherwise be used by ATP synthases for ATP production. Assuming a maximal H^+^/ATP ratio of 4 [28], the total energetic cost of the Pgl bypass is 1.25 ATP equivalents per molecule of glucose entering the oxPPP, corresponding to an extra ATP demand of 1.33 mmol·[g_CDW_ · h]^−1^. In other words, roughly 20 % of the energy used by the cell for glucose uptake and subsequent phosphorylation by the PTS system is wasted in the Pgl bypass. Furthermore, because gluconolactone hydrolysis occurs non-enzymatically outside the cell, it escapes direct metabolic control and the gluconolactone and gluconate are potentially accessible to other cells competing for carbon sources. An illustrative example of this kind of cross-feeding is the recovery of normal growth in Δ*zwf*Δ*pgi E. coli* in the presence of Δ*pgl* mutants, likely because the gluconolactone or gluconate released by the latter feed the oxPPP of the Δ*zwf*Δ*pgi* cells [15]. The downsides of this extracellular bypass are therefore that it is more energetically expensive, partially beyond metabolic regulation and subject to hijack by competing cells.

Two oxPPP topologies must therefore be considered in metabolic flux analyses depending on whether Pgl is present (the canonical oxPPP) or absent (with Pgl bypass). The latter configuration is generally ignored however [7,29], particularly in analyses of *E. coli* BL21 [30,31], which is widely used in biotechnological applications, notably for heterologous protein production [32]. *E. coli* BL21 and derived strains lack the *pgl* gene and there is evidence of Pgl bypass activity in this strain [33].

Our results show that metabolic activity in the Pgl bypass oxPPP is similar to that of the canonical oxPPP in the WT strain. As observed previously though [7,14,15], the Δ*pgl* mutant grows about 30% slower than the WT. The growth rate is reduced even further if gluconate utilisation is also blocked (down to about 50% of the WT’s growth rate), highlighting the metabolic contribution of the Pgl bypass. Finally, of the three enzymes involved in the oxPPP (Pgl, G6P dehydrogenase, and 6PGNT dehydrogenase, the latter two being encoded by *zwf* and *gnd* respectively), Pgl is the only one whose absence is associated with a noticeable growth defect. In particular, the silent growth phenotype of the Δ*zwf* mutant demonstrates that flux in the oxPPP is not absolutely required for bacterial growth on glucose. Deletion of *zwf* leads to a global reorganization of metabolic fluxes through *E. coli*’s central metabolism to compensate for the absence of flux in the oxPPP while providing the required precursors and energy for anabolism [6,34,35]. In contrast, *pgl* deletion leads to a local rearrangement of carbon fluxes that fails to support optimal growth. Furthermore, in spite of the secretion mechanism described in this study, absence of Pgl must nevertheless lead to intracellular accumulation of δ-6PGL, Pgl’s substrate, and γ-6PGL, which forms faster by isomerization of δ-6PGL than δ-6PGL spontaneously hydrolyses to 6PGNT [11]. Indeed, δ-6PGL accumulation has been observed in *E. coli* BL21 growing on glucose [33]. The accumulation of these reactive compounds may be detrimental to cells. The growth defect of the Δ*pgl* mutant can therefore be explained by the energy cost of the Pgl bypass and/or intracellular accumulation of toxic intermediates.

Pgl is often described as predominantly a house-cleaning enzyme, preventing the formation of undesirable by-products [9,12]. According to this view, oxPPP flux should mainly be maintained by non-enzymatic hydrolysis of 6PGL into 6PGNT [7,12,14]. Our results shown on the contrary that the contribution of non-enzymatic δ-6PGL hydrolysis to the canonical oxPPP in the absence of Pgl is negligible, a situation (absence of Pgl) that should promote non-enzymatic hydrolysis of δ-6PGL, since the intracellular concentration of the latter is increased, as mentioned above [33]. We can therefore assume that non-enzymatic hydrolysis of δ-6PGL is also negligible when Pgl is present (*i*.*e*. in the WT strain), meaning that the Pgl’s catalytic activity is crucial for 6PG formation. Our results therefore suggest that Pgl’s catalytic role is crucial for the efficient functioning of the central metabolic network. While Pgl does also help prevent intracellular accumulation of 6PGLs, our results suggest that this role is mediated by an additional process involving the excretion of these compounds from the cell.

In summary, our comprehensive functional analysis of a knock-out mutant in *E. coli*’s central metabolism confirms the existence of a long described compensatory mechanisms and reveals fundamental gaps in our current understanding of an enzyme operating in parallel to a non-enzymatic reaction. Because the reaction catalysed by Pgl can occur spontaneously, the role of Pgl in metabolism has remained unclear. Our results indicate that Pgl’s catalysis of 6PGL hydrolysis into 6PGNT is crucial to maintain the required flux through the oxPPP in an energy-efficient manner.

## METHODS

### Strains and media

*Escherichia coli* BW25113 K-12 was selected as the experimental model (wild-type strain) for this study. *E. coli* BW25113 *Δpgl* was constructed from a Δ*pgl* strain in the Keio collection [36], with kanamycin cassette removal with a pCP20 plasmid encoding FLP recombinase [37]. The deletion mutant *ΔpglΔgnt* was obtained via a one-step disruption protocol [38], and the triple deletion mutant *ΔpglΔgntΔIdnK* similarly from the double *ΔpglΔgnt* mutant. To confirm the mutations, polymerase chain reaction (PCR) was used to amplify fragments containing the modified sequences. The lengths of the amplified fragments were tested by agarose gel electrophoresis and compared with those of the previous mutant strain. The complemented strain (*Δpgl::pgl*) was obtained by transformation of the *Δpg*l strain by a pZA23::*pgl* plasmid carrying the *pgl* gene under the control of the pLac promoter. The *pgl* gene was amplified from the *E. coli* BW25113 WT strain by PCR and inserted in pZA23 plasmid using the In-Fusion® HD Cloning Plus CE Kit (Takara).

All *E. coli* strains were grown in M9-based minimal synthetic medium as described in [6], complemented with 15 mM glucose. Cultures (50 mL) were performed in triplicate in baffled shake flasks at 37 °C and 200 rpm. Growth was monitored by measuring the optical density at 600 nm (OD_600_) using a Genesys 6 spectrophotometer (Thermo, USA), and a correlation factor of 0.37 g_CDW_·(L·OD_600_)^−1^ was used to calculate biomass concentration.

### NMR analysis

NMR spectra were recorded on a Bruker Avance III HD 800 MHz spectrometer equipped with a 5-mm quadruple-resonance QCI-P (H/P-C/N/D) cryogenically cooled probe head. D_2_O (10 vol.%) was added to the samples for field/frequency locking and 1 mM TSP-d4 (dissolved in D_2_O) was added as an internal standard for frequency calibration and concentration measurements. Spectra were recorded and processed using Bruker TopSpin 3.2. 1D ^1^H NMR spectra were acquired using the zgpr30 sequence at 280 K with 32 or 64 scans, 64k points, an acquisition time of 2 s, and an recycle delay of 8 s.

### Extracellular metabolite concentration measurements and calculation of extracellular fluxes

Metabolite concentrations (glucose, acetate, gluconate) were quantified by 1D ^1^H NMR from the supernatant obtained by centrifugation (12,000 g, 3 min) of culture broth. The 1D ^1^H NMR data were acquired as described above (64 scans). For samples containing gluconolactone, the metabolites (including glucose, gluconate and gluconolactone) were quantified immediately after filtration (0.2 µm, polyethersulfone membrane) of 500 µL of culture. The filtrate was kept on ice and rapidly analysed by NMR as described above (32 scans).

Extracellular fluxes (*i*.*e*. glucose uptake, GNT accumulation and growth rates) were determined from the time course concentrations of biomass, substrates, and products using PhysioFit v2.0.4 [39](https://github.com/MetaSys-LISBP/PhysioFit).

### Stationary ^13^C-labelling experiments

After preculture on LB, strains were grown in 50 mL M9 minimal synthetic medium complemented with 15 mM [1-^13^C] glucose in baffled shake flasks at 37 °C and 200 rpm. At OD 1.2, cells were harvested by centrifugation (5 min at 10,000 g) and the pellet was resuspended in 1.250 mL of Milli-Q H_2_O. The suspension (250 µL) was mixed with 250 µL of HCl 12 N and hydrolysed at 110 °C for 18 h. HCl was evaporated using a vacuum concentrator, the pellet was washed twice with Milli-Q H_2_O and resuspended in 100 µL H_2_O. This sample was diluted 100 times for analysis. The carbon isotopologue distribution of alanine was measured in three independent biological replicates, as detailed in [40].

### ^13^C-Pulse experiments

*E. coli* BW25113 *Δpgl* was grown in 50 mL M9 minimal synthetic medium complemented with 15 mM glucose in a baffled shake flask at 37 °C and 200 rpm. At OD 1.2, [2-^13^C] glucose was added to the culture medium to obtain approximately 50 % of ^13^C-labelled glucose. Growth was monitored using OD_600_ measurements as described above. Extracellular compounds were quantified directly by NMR as described above.

### Kinetics of phosphogluconolactone and gluconolactone spontaneous hydrolysis

*Real time monitoring of gluconolactone (GL) degradation*. δ-Gluconolactone (δ-GL) degradation was monitored by ^1^H NMR using a pseudo-2D pulse program (noesyphpr) at 310K. A 3 mM δ-GL (Sigma-Aldrich) solution was prepared in M9-based minimal synthetic medium. Acquisitions were started after rapid homogenisation and temperature stabilisation, and spectra were recorded every 65 s with 8 scans each (acquisition time, 3 s; recycle delay, 5 s) for a total of 120 time points. δ-GL degradation at 280 K was monitored using the same procedure and a freshly prepared δ-GL solution.

*Real time monitoring of 6-phosphogluconolactone (6PGL) degradation*. 6PGL was produced enzymatically from glucose-6-phosphate (G6P) using commercial *L. mesenteroides* NADP^+^ dependent G6P-dehydrogenase (G6PDH) expressed in *E. coli* (Sigma-Aldrich). The reaction mix consisted of 5 mM G6P, 5 mM NADP^+^, 12 mM MgCl_2_, 100 mM phosphate buffer (at pH = 7.2) and 1 mM TSP-d4 [6]. ^1^H and ^31^P 1D NMR spectra of the mix were recorded using the zgpr30 sequence with 16 scans and the zg sequence with 64 scans, respectively. One enzyme unit of G6PDH was then added to the NMR tube and the production and hydrolysis of 6PGL were monitored at 310 K using dual reception (^1^H and ^31^P) pseudo-2D spectra (2DDR zggpw5) [41]. Spectra were recorded every 115 s with 64 scans each for a total of 64 time points (^1^H acquisition time, 0.7 s; recycle delay; ^31^P acquisition time 0.6 s, recycle delay, 1 s). The concentrations of G6P, 6PGL and 6PGNT were determined from the rows of the pseudo-2D ^31^P spectra.

*Calculation of degradation constants*. The degradation constants of GL and 6PGL were determined by fitting their time-course concentrations assuming a first-order process using COPASI [42] (v4.27), as detailed in [17].

### Relative contributions of glycolysis and the oxPPP

The contributions of the oxPPP and Emben-Meyerhof-Parnas (EMP) pathway to glucose metabolism were estimated by quantifying alanine isotopologues. [1-^13^C] glucose metabolised through the EMP pathway forms unlabelled and [1-^13^C] pyruvate in equal proportions, while C_1_-decarboxylation of glucose through the oxPPP only produces unlabelled pyruvate. The contributions of the oxPPP and the EMP pathways were thus estimated for each biological replicate from the fraction of the M1 isotopologue of alanine, using the following algebraic equations [21]:

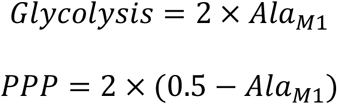

The mean and standard deviation (SD) of the relative contributions of glycolysis and oxPPP were calculated using the values determined from three independent biological replicates (n=3).

### Dynamic ^13^C-flux model of glucose metabolism to quantify the contribution of the Pgl bypass

To quantify the contributions of the canonical oxPPP and of the Pgl bypass to ribose-5-phosphate biosynthesis, we constructed a dynamic ^13^C-flux model of glucose metabolism following the formalism detailed in [43]. The model contains 21 reactions, 25 species, and 2 compartments (the environment and the cell), and represents five processes: i) growth, ii) glucose uptake and phosphorylation into G6P, utilisation of G6P through iii) the EMP pathway and iv) the intracellular branch of the oxPPP, and v) the extracellular Pgl bypass (Figure 7).

The differential equations, which balance the concentrations of extracellular compounds (biomass, glucose, gluconolactone and gluconate) and intracellular compounds (G6P, glyceraldehyde-3-phosphate, 6PGL, 6PGNT and pyruvate), were completed with isotopic equations for parameter estimation. As detailed in [43,44], we considered all reactions (except biomass synthesis) separately for unlabelled and labelled reactants. Fluxes were assumed to be constant over time, in line with the metabolic steady-state assumption, except for the gluconolactone hydrolysis flux which was modelled using the mass action law to represent first-order degradation kinetics [17], in keeping with the abiotic hydrolysis of gluconolactone into gluconate (see Results).

The final model has 8 free parameters in total (Supplementary material). The contribution of the oxPPP to glucose metabolism (*i*.*e*. the sum of the flux through the intra- and extracellular branches of the oxPPP) and the gluconolactone (abiotic) hydrolysis rate were fixed at the experimental values determined in this study, as detailed in the Results section. The remaining parameters (p) were estimated by fitting to the experimentally determined concentration dynamics of biomass and unlabelled and labelled glucose and gluconolactone, by minimising the objective function *f* defined as the weighted sum of squared errors:

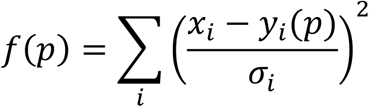

where *x*_*i*_ is the experimental value of data point *i*, with an experimental standard deviation *σ*_*i*_, and *y*_*i*_*(p)* is the corresponding simulated value. The objective function *f* was minimised using the particle swarm optimisation algorithm (2,000 iterations with a swarm size of 50). The experimental and fitted data of one biological replicate are shown in Figure 7, and the results for all replicates are provided in the Supplementary material.

The model was constructed and analysed using COPASI [42] (v4.27) and is provided in SBML and COPASI formats in the Supplementary material and at https://github.com/MetaSys-LISBP/GL_GNT_bypass with the corresponding data for all replicates.

## CRediT authorship contribution statement

**LP**: Conceptualization, Formal analysis, Investigation, Writing-Original draft, Writing-Reviewing and Editing, Visualization. **JP**: Conceptualization, Formal analysis, Investigation, Writing-Original draft. **SUJ**: Conceptualization, Formal analysis, Investigation, Writing-Original draft, Writing-Reviewing and Editing, Visualization. **EC**: Methodology. **PM**: Conceptualization, Software, Formal analysis, Investigation, Data curation, Writing-Original draft, Writing-Reviewing and Editing, Visualization. **FL**: Conceptualization, Validation, Data curation, Writing-Original draft, Writing-Reviewing and Editing, Supervision, Funding acquisition.

## Data availability

Dataset was deposited into a publicly accessible repository (https://data.mendeley.com/preview/2rw5pz75yr).

The dynamic ^13^C-flux model has been deposited in the Biomodels database (https://www.ebi.ac.uk/biomodels) [45] with the identifier MODEL2310250001.

## Supporting information

This article contains supporting information.

Supplementary material: Model documentation

## Acknowledgements

This study was funded by the Agence Nationale de la Recherche, grant number ANR-19-CE44-0005 (PerioMet).

The authors thank MetaToul (Metabolomics & Fluxomics Facilities, Toulouse, France, www.metatoul.fr) and its staff for technical support and access to the NMR facility. MetaToul is part of the French National Infrastructure for Metabolomics and Fluxomics (www.metabohub.fr), funded by the ANR (MetaboHUB-ANR-11-INBS-0010).

## 1. Model overview

This model contains a coarse-grained representation of the central metabolism and of the extracellular gluconolactone-gluconate (GL-GNT) by-pass of *Escherichia coli*.

This model was developed with COPASI (http://copasi.org). All models are available in SBML and COPASI formats at https://github.com/MetaSys-LISBP/GL_GNT_bypass. The model can also be downloaded from the Biomodels database (http://www.ebi.ac.uk/biomodels) with identifier MODEL2310250001.

This model comprises 2 compartments (the environment and the cell), 9 species (8 metabolites and biomass) and 9 reactions that represent the following processes (Figure 1):

▪ glucose uptake
▪ glycolysis
▪ pentose phosphate pathway
▪ extracellular GL-GNT by-pass
▪ growth

**Figure 1.**
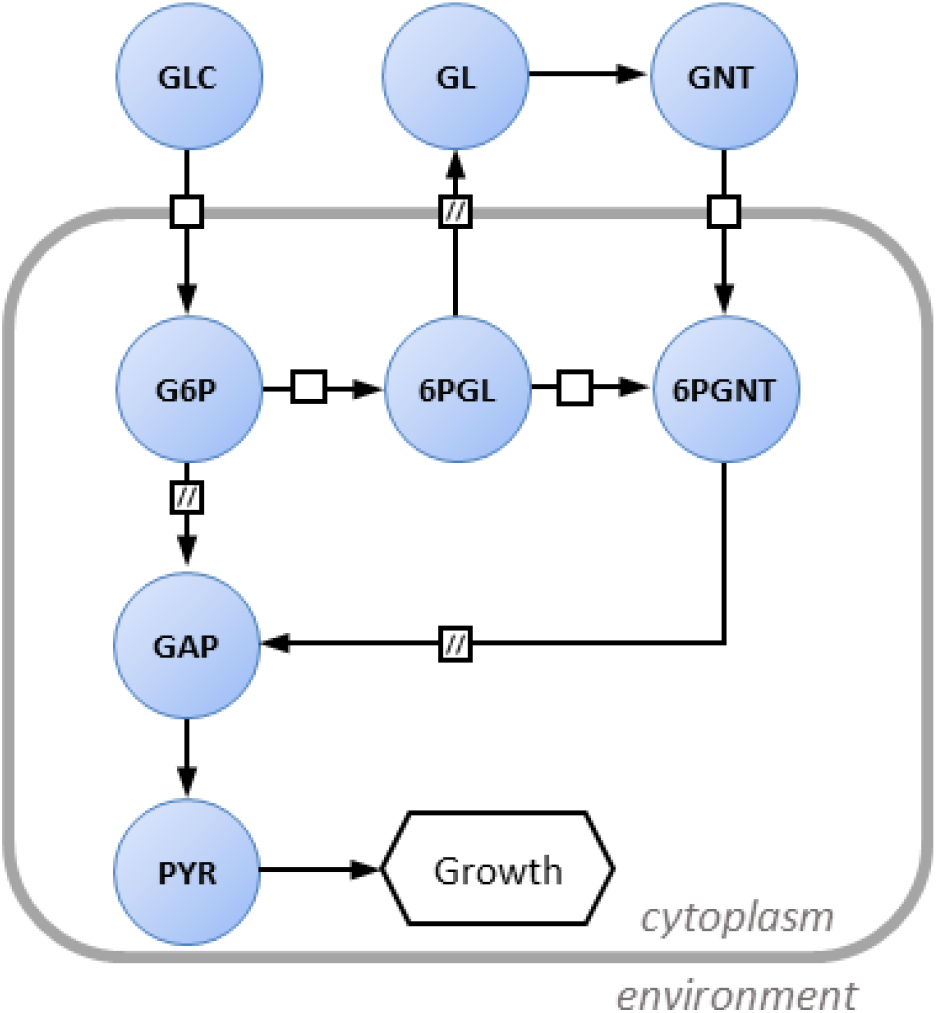
Metabolic network of *E. coli* metabolism. The diagram follows the conventions of the Systems Biology Graphical Notation process description.

## 2. Model units

Model units are millimole (mmol) for amounts of metabolites, litre (L) for volumes, and hour (h) for time. Amount of biomass is expressed as gram dry weight (g_DW_).

## 3. Reactions

The reactions included in the model are listed in the table below.

| *Name* | *Reaction* | *Flux* |
| --- | --- | --- |
| r1_PTS | GLC → G6P | v1 |
| r2_EMP_high | G6P → GAP | v2 |
| r3_EMP_low | GAP → PYR | v3 |
| r4_PYR_out | PYR → ∅ | v4 |
| r5_G6PDH | G6P → 6PGL | v5 |
| r6_PGL | 6PGL → 6PG | v6 |
| r7_PPP | 6PG → 1.66 * GAP | v7 |
| r8_GL_export | 6PGL → GL<sub>env</sub> | v8 |
| r9_GL_degradation | GL<sub>env</sub> → GNT<sub>env</sub> | v9 |
| r10_GNT_uptake | GNT<sub>env</sub> → 6PG | v10 |
| r11_growth | X → 2 * X | v11 |

## 4. ODEs system

The differential equations, which describe the progression of the variables over time as a function of the system’s rates, balance the concentrations of extracellular (biomass, glucose, gluconolactone and gluconate) and intracellular (G6P, 6PGL, 6PG, GAP and PYR) species.

Extracellular species:

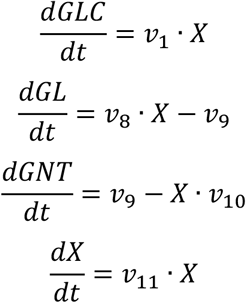

Intracellular species:

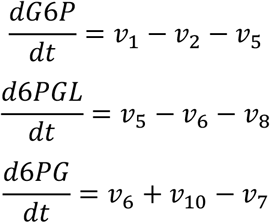

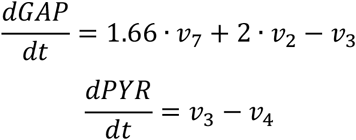

## 5. Reaction rates

Non-enzymatic degradation of extracellular gluconolactone was modelled using a first-order degradation rate law (*v*_9_ = *k*_*deg* ·_ [*GL*]), as observed experimentally. Glucose uptake rate (*v*_1_) and growth rate (*v*_11_) were defined as constant. All other reaction rates were constrained to match the steady-state conditions (exponential growth) and balance all intracellular species. The following constraints were used:

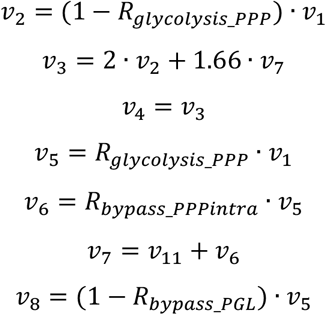

where *R*_*glycolysis_PPP*_ is the flux partition between glycolysis and the oxidative pentose phosphate pathway (i.e.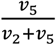) and *R*_*bypass_PGL*_ is the flux partition between PGL and the extracellular GL-GNT bypass (i.e.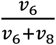).

Since no accumulation of gluconate was detected, we also balanced its concentration:

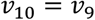

## 6. Extension with isotopic equations

This dynamic model was extended with isotopic equations as detailed previously (doi: 10.1186/s12918-015-0213-8). Briefly, all reactions (except biomass synthesis) were considered separately for unlabeled and labeled metabolites. For instance, the isotopically extended balance of G6P corresponds to:

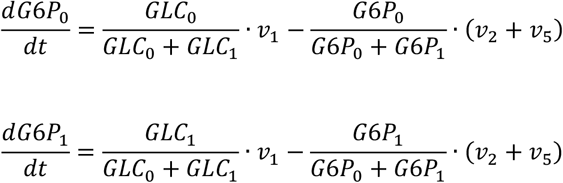

where subscripts 0 and 1 refers to the unlabeled and labeled metabolite, respectively.

## 7. Pulse of ^13^C-glucose

To simulate the isotopic dynamics in response to a pulse of ^13^C-labeled glucose, we added an event which sets the concentration of ^13^C-glucose (*GLC*_1_) at time *t*_*pulse*_.

## 8. Flux calculation

Initial concentrations of unlabelled intracellular metabolites were set to unity, and initial concentrations of labelled metabolites were set to zero. The degradation constant of GL was set to its experimental value (*k*_*deg*_ = 2.58 *h*^−1^). The flux partition between glycolysis and the oxidative pentose phosphate pathway was set to its experimental value (*R*_*glycolysis_PPP*_ = 0.17, calculated by summing the relative contribution of the PPP to PYR biosynthesis determined by stationary ^13^C-labeling experiments – 0.15 – and the accumulation rate of extracellular GL relative to glucose uptake rate – 0.17/8.52 = 0.02).

This model therefore contains 7 free parameters:

- initial concentrations of extracellular glucose (*GLC*_0_) and biomass (*X*)
- time at which the pulse of ^13^C-glucose is performed (*t*_*pulse*_)
- concentration of ^13^C-glucose added during the pulse (*GLC*_1_)
- glucose uptake rate (*v*_1_) and growth rate (*v*_11_)
- flux partition between the extracellular GL-GNT by-pass and PGL (*R*_*bypass_PGL*_).

These parameters were estimated by fitting experimental data (time course concentrations of unlabelled and labelled glucose and gluconolactone, and time course concentration of biomass), as detailed in the publication. Values and standard deviations obtained for each of the three independent biological replicates are provided below. All parameters could be determined with a good precision in each biological replicate. We recalculated all fluxes from the estimated parameters.

| **Parameter** | **Replicate 1** |  | **Replicate 2** |  | **Replicate 3** |  | **mean<sup>c</sup>** | **sd<sup>c</sup>** |
| --- | --- | --- | --- | --- | --- | --- | --- | --- |
|  | *value<sup>a</sup>* | *sd<sup>b</sup>* | *value<sup>a</sup>* | *sd<sup>b</sup>* | *value<sup>a</sup>* | *sd<sup>b</sup>* |  |  |
| $GLC_0$ | 15.59 | 0.13 | 15.00 | 0.10 | 15.04 | 0.09 | **15.21** | **0.33** |
| $X$ | 0.057 | 0.003 | 0.052 | 0.002 | 0.051 | 0.002 | **0.053** | **0.004** |
| $GLC_1$ | 12.2 | 0.2 | 9.3 | 0.2 | 10.2 | 0.2 | **10.6** | **1.5** |
| $t_{pulse}$ | 5.2 | 0.1 | 5.0 | 0.1 | 5.0 | 0.1 | **5.0** | **0.1** |
| $R_{bypass\_PGL}$ | 0.001 | 0.027 | 0.001 | 0.026 | 0.001 | 0.026 | **0.001** | **0.001** |
| $v_1$ | 7.75 | 0.15 | 7.04 | 0.12 | 7.10 | 0.13 | **7.29** | **0.39** |
| $v_{11}$ | 0.395 | 0.007 | 0.412 | 0.006 | 0.410 | 0.006 | **0.406** | **0.009** |

<sup>a</sup>best fit obtained with COPASI
<sup>b</sup>standard deviation estimated by COPASI from the best fit
<sup>c</sup>mean and standard deviation of the three independent biological replicates

## Abbreviations

6PGL: 6-phosphogluconolactone
6PGNT: 6-phosphogluconate
δ-6PGL: δ-6-phosphogluconolactone
DHO: dihydroorotate
GNT: gluconate
EMP: Embden-Meyerhof-Parnas
GAP: glyceraldehyde-3-phosphate
GL: gluconolactone
GLC: glucose
G6P: glucose-6-phosphate
GntK: gluconate kinase
IdnK: thermosensitive gluconate kinase
NMR: nuclear magnetic resonance
oxPPP: oxidative branch of the pentose-phosphate pathway
Pi: inorganic phosphate
PTS: phosphoenolpyruvate:glucose phosphotransferase system
R5P: ribulose-5-phosphate
WT: wild-type
Zwf: glucose-6-phosphate dehydrogenase

