## Supplementary_Data for "6-Phosphogluconolactonase is critical for the efficient functioning of the pentose phosphate pathway" (from another version of this preprint: version 3, 2024-04-08)

-

**SUPPLEMENTARY MATERIAL 1**

Léa Phégnon[1‡], Julien Pérochon[1‡], Sandrine Uttenweiler-Joseph[1], Edern Cahoreau[1,2], Pierre Millard[1,2] and Fabien Létisse[1,3*]

[1]Toulouse Biotechnology Institute, Université de Toulouse, INSA, UPS, Toulouse, France.

[2]MetaToul-MetaboHUB, National Infrastructure of Metabolomics and Fluxomics, Toulouse, France.

[3]Institut de Pharmacologie et de Biologie Structurale (IPBS), Université de Toulouse, CNRS, Université Toulouse III - Paul Sabatier (UT3), Toulouse, France

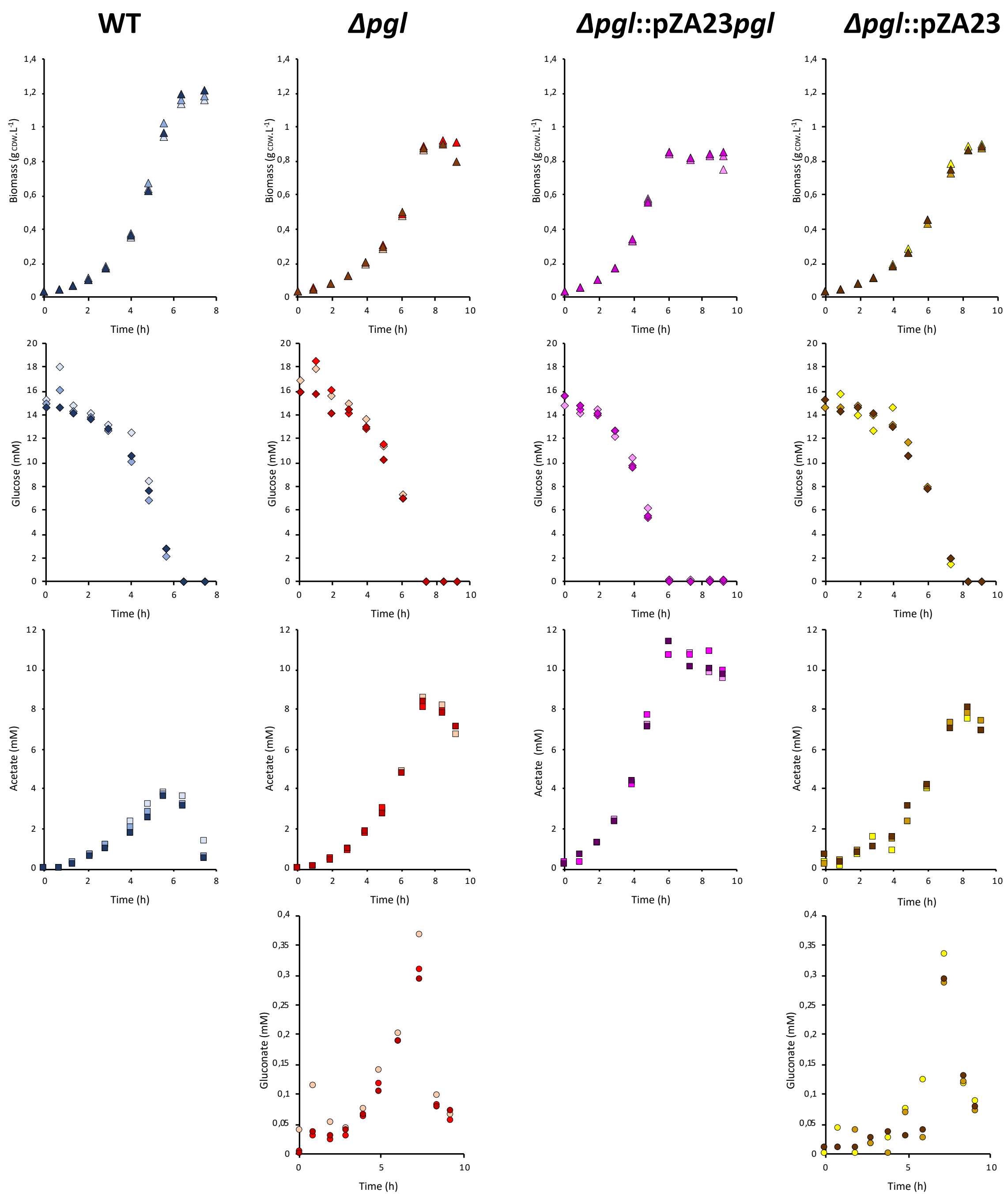


**Supplementary figure 1. Biomass (Δ) and extracellular metabolite (glucose (◊), acetate (□) and gluconate (○)) profiles of cultures (n=3 biological replicates, colour gradient) of *E. coli* BW25113 WT (blue), and *Δpgl* (red), *Δpgl*::pZA23*pgl* complement (purple) and *Δpgl*::pZA23 control (yellow) strains.** Cultures were performed in M9-based medium supplemented with glucose at pH 7. Note that gluconate was only detected in the *Δpgl* and the *Δpgl*::pZA23 strains.

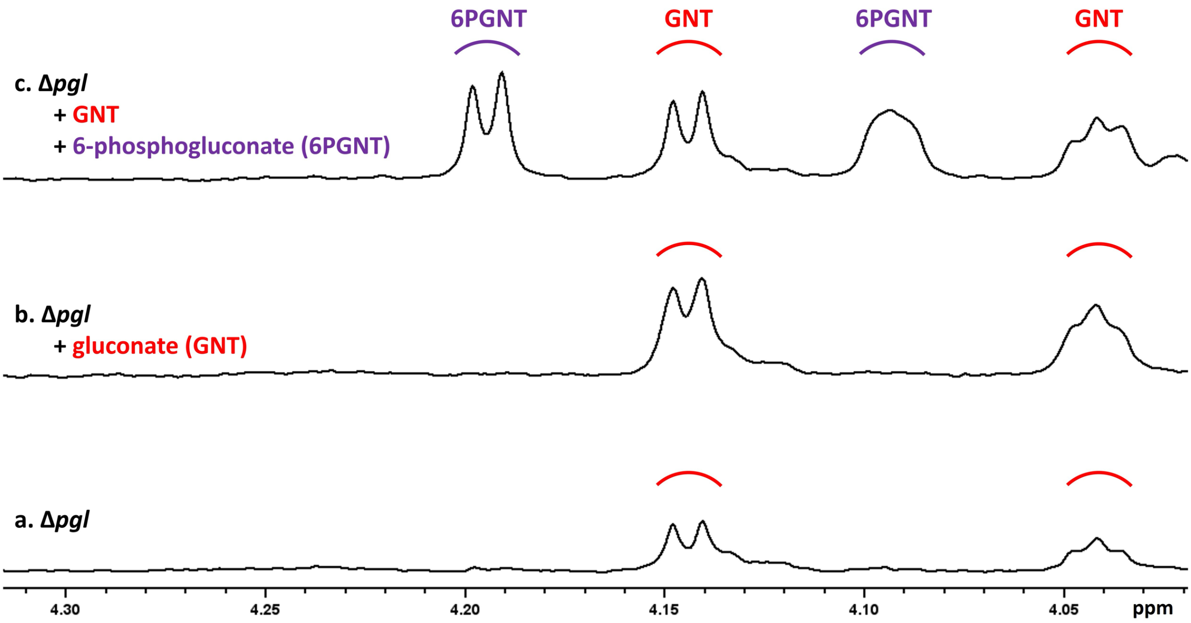


**Supplementary figure 2. Gluconate identification by $^{1}$H NMR analysis of *E. coli* BW25113 Δ*pgl* culture supernatants.**

Selected region of the 1D $^{1}$H NMR spectrum of the culture supernatant of **a.** *E. coli* BW25113 Δ*pgl* (estimated concentration, 0.35mM), **b.** *E. coli* BW25113 Δ*pgl* spiked with 0.2mM gluconate and **c**. *E. coli* BW25113 Δ*pgl* after spiking of 6-phosphogluconate (6PGNT) at 0.7mM.

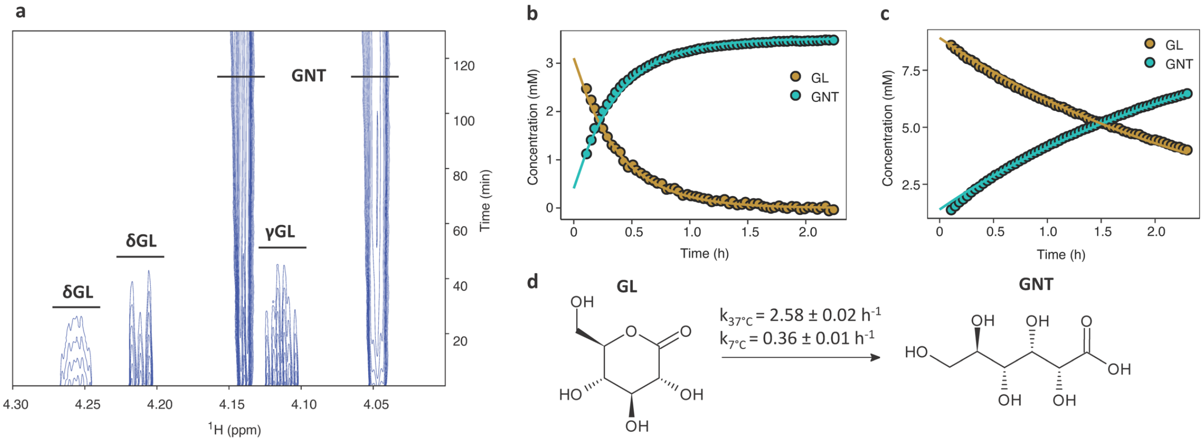


**Supplementary figure 3. Gluconolactone spontaneous hydrolysis**. **a**. Pseudo-2D $^1$H spectra at 37 °C, pH 7.2 in M9 synthetic minimal media of spontaneous gluconolactone hydrolysis. T0 corresponds to the beginning of NMR acquisition (= 6 min 35 s after gluconolactone addition). The signals at 4.05 and 4.14 ppm correspond respectively to the protons attached to the carbon-3 (C3) and -2 (C2) of the gluconate. **b,c**. GL and GNT concentrations extracted from pseudo-2D spectra (dots) and fitted by COPASI model (lines) at 37 °C (**b**) and 7 °C (**c**). **d.** Reaction scheme for the spontaneous hydrolysis of gluconolactone (GL) into gluconic acid (GNT) with the degradation constants obtained at 37 °C and 7 °C at pH 7.2.

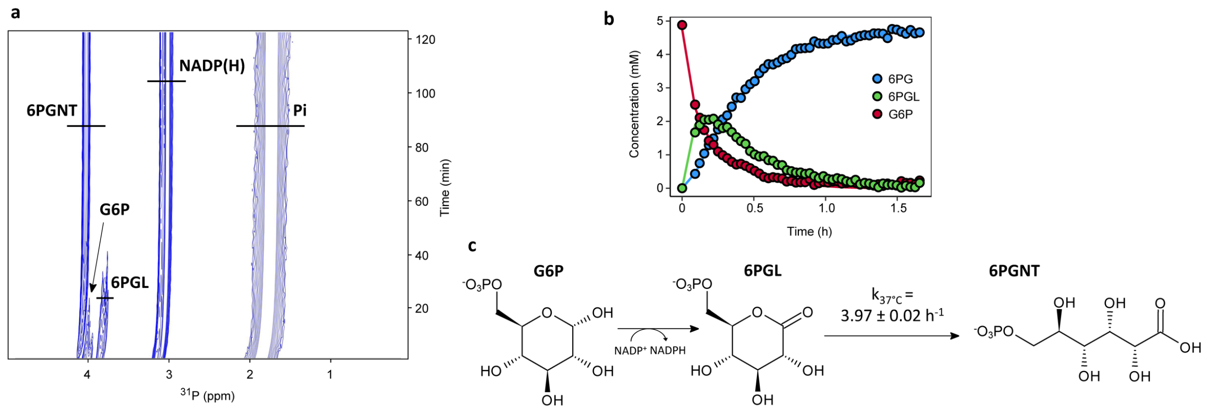


**Supplementary figure 4: Enzymatic production and spontaneous hydrolysis of 6-phosphogluconolactone (6PGL) at 37 °C**. **a**. Pseudo-2D $^{31}P$ spectra of 6-phosphogluconolactone (6PGL) recorded at 37°C, pH 7.2 in phosphate buffer. T0 corresponds to the start of NMR acquisition (5 min 25 s after 1U glucose-6-phosphate dehydrogenase (G6PDH) addition). 6PGNT, 6-phosphogluconate; G6P, glucose-6-phosphate; 6PGL, 6-phosphogluconolactone; Pi, inorganic phosphate. **b**. Concentrations of 6-phosphogluconate (6PG), glucose-6-phosphate (G6P) and 6PGL (dots) extracted from the pseudo-2D spectra and fitted using a COPASI model (lines). **c**. Reaction scheme of enzymatic production of 6PGL from G6P via G6PDH and spontaneous 6PGL hydrolysis into 6PGNT, with the value of the degradation constant (k) obtained at 37 °C and at pH 7.2.

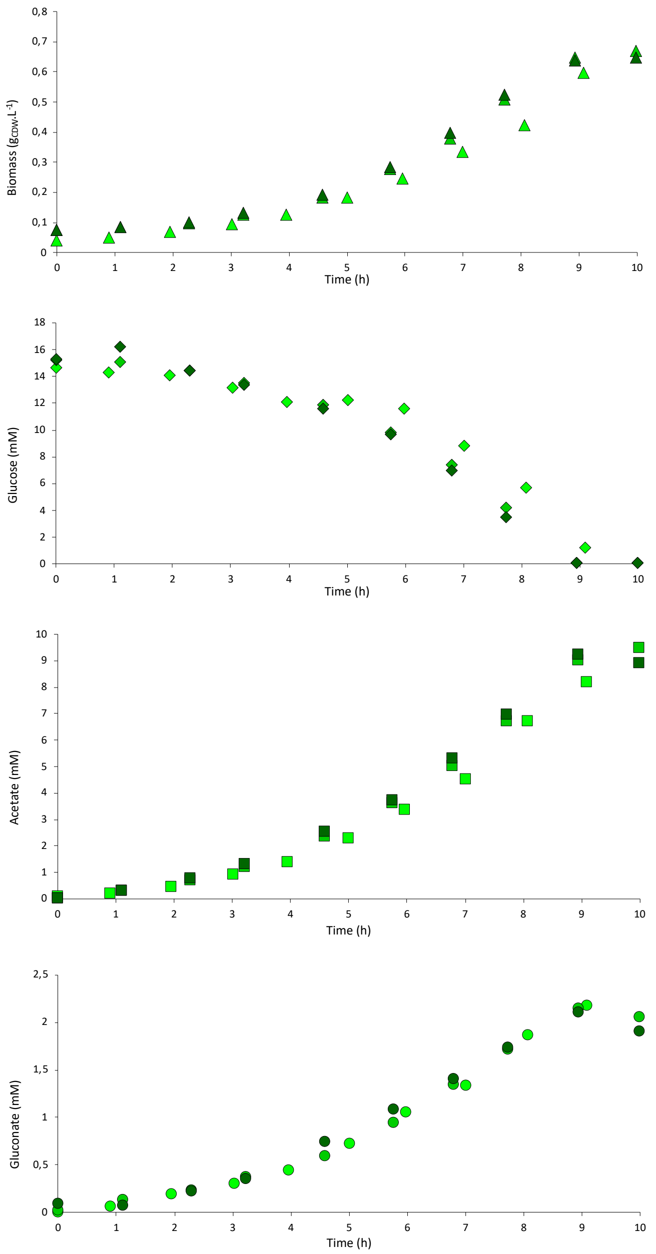


**Supplementary figure 5: Biomass (Δ) and extracellular metabolite (glucose (◊), acetate (□) and gluconate (○)) profiles of cultures (n=3 biological replicates, colour gradient) of *E. coli* BW25113 Δ*pgl*Δ*gntK*Δ*idnK*.** Cultures were performed in M9-based medium supplemented with glucose at pH 7.
